# Galectin-14 expression in ovarian cancer

**DOI:** 10.1101/717793

**Authors:** Lorenna Oliveira Fernandes de Araujo, Yves St-Pierre

**Author notes:** **Corresponding author:** Yves St-Pierre, INRS-Institut Armand-Frappier, 531 Boul. Des Prairies, Laval, Québec, Canada, H7V 1B7. Abbreviations: galectin-14 (GAL-14), ovarian cancer (OC).

## Abstract

Galectins (gal) are multifunctional proteins whose expression changes under different physiological or pathological conditions, including cancer. However, so far, most studies have focused on gal-1 and gal-3, and to a lesser extent to gal-7 and gal-9. We still know very little about other galectins, especially the recently discovered ones, such as gal-14, a prototype galectin highly expressed at the maternal-fetal interface. Here, using *in silico* and *in vitro* approaches, we report a correlation between *lgals14* expression and ovarian cancer. We found that high expression of gal-14 mRNA in ovarian cancer cells is associated with a shorter survival. Consistent with this observation, we also found that *lgals14* is preferentially expressed in high grade serous adenocarcinoma (HGSA) ovarian cancer. Our *in vitro* data with ovarian cancer cell lines confirmed that *lgals14* is readily expressed in HGSA. Interestingly, *de novo* expression of gal-14 in HEK-293 cells increased apoptosis, both at the basal level and following exposure to low doses of etoposide. Thus, although the study of this galectin is still in its infancy, we were able to provide novel insights into the expression patterns of this galectin and its involvement in cancer.

## Introduction

Galectins are small molecular weight soluble proteins with have high affinity for β-galactosides [1]. They were first recognized as lactose-specific lectins, or more specifically β-galactoside-binding lectins, a group of soluble proteins widely found in extracts of vertebrate tissues and whose activity could be inhibited by lactose, galactose or related carbohydrates, such as *N-*acetylgalactosamine [2–4]. To date, in mammals, a total of 19 different galectins have been identified. They are involved in a large number of cellular functions, most notably in the regulation of cell survival. In vertebrates, galectins are found in a variety of tissues and in almost every cell type. Gal-1, for example, is widely expressed by various cell types, such as vascular endothelial cells, epithelial cells and cells from both primary and secondary lymphoid organs [5–7]. Gal-3, another extensively documented galectin, is expressed in a variety of human tissues and cells, such as in normal human peripheral blood monocytes, activated macrophages, mast cells, T cells, certain epithelial cells and sensory neurons [8–10]. Gal-7 is largely expressed in normal epithelial cells and is often considered a marker of epithelial differentiation [11, 12]. The two tandem-repeat galectins, gal-8 and −9, are expressed in vascular and lymphatic endothelial cells, and in the thymus, respectively [13, 14]. The Charcot-Leyden Crystal protein, or gal-10, is abundantly expressed in the bone marrow, the birthplace of immune cells, and in several subpopulations of immune cells [5, 15, 16]. The expression pattern and role of of other galectins is, however, less well known. We do know, however, that gal-12 is predominantly expressed in adipocytes while gal-13, −14 and −16 are preferentially expressed in the placenta [16–19].

Gal-14, also known as Placental Protein 13-like or PPL 13, is one the less well known galectins. Gal-14 is a prototype galectin with significant homology to gal-13 [17]. Galectin-14 gene is identified as *lgals14* and located within a cluster of galectin genes on the q13.2 region of chromosome 19 [16, 17]. Although it has been mostly known for its expression in the placenta, studies with animal models have shown that gal-14 is also expressed in eosinophils from sheep, being released from these cells in response to allergen stimulation, suggesting a role of gal-14 in the regulation of eosinophils activity during allergic responses [20]. Its expression and role in cancer, however, remains largely unknown. In the present work, we provide novel insights into the expression patterns of this galectin and explore its potential role in regulating cell survival in cancer.

## Results

### In silico analysis of galectin-14 expression in normal and cancer tissues

To obtain a global view of gal-14 expression in normal and cancer tissues, we first interrogated TCGA datasets. As expected, the tissue expressing the highest number of *lgals14* transcripts was the placenta as compared with other normal tissues (Figure 1). In cancer tissues, *lgals14* was expressed at a higher level in several types of cancer, including liver, breast, liver, uterine and ovarian cancer (Figure 2a). In contrast, levels of *gals14* were relatively low in prostate, uveal, and pancreas cancer, indicating that *lgals14* expression in cancer is cancer-type specific. Analysis of *lgals14* expression in the NCI-60 panel of human cancer cell lines that is commonly used to assess the expression of specific genes in cancer confirmed that *lgals14* expression is significantly higher in human ovarian cancer cell lines when compared to other cancer cell lines (Figure 2b).

**Figure 1:**
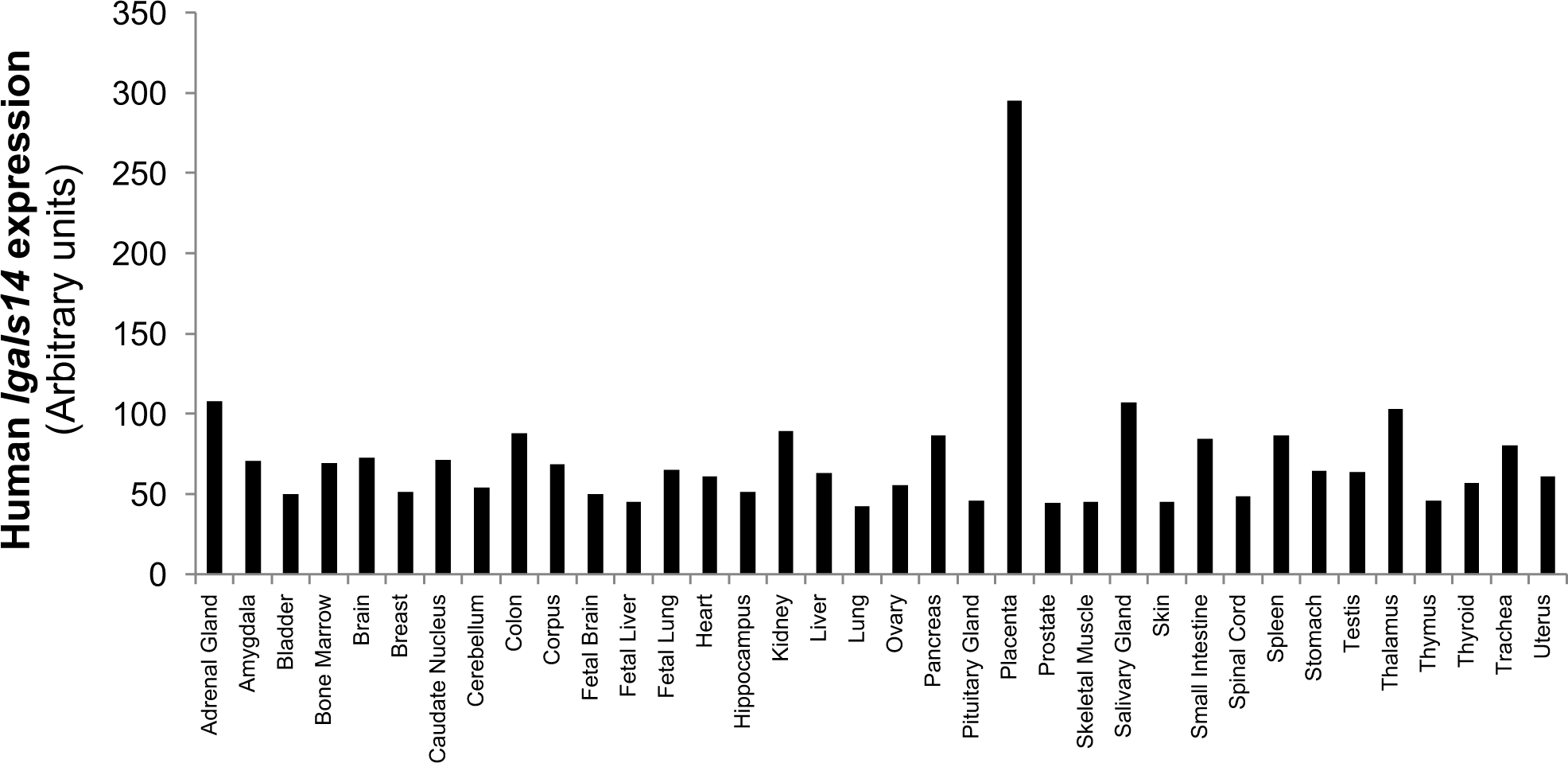
Expression of lgals14 in normal human tissues. Data were generated using the publicly availabe GeoProfile databas.

**Figure 2:**
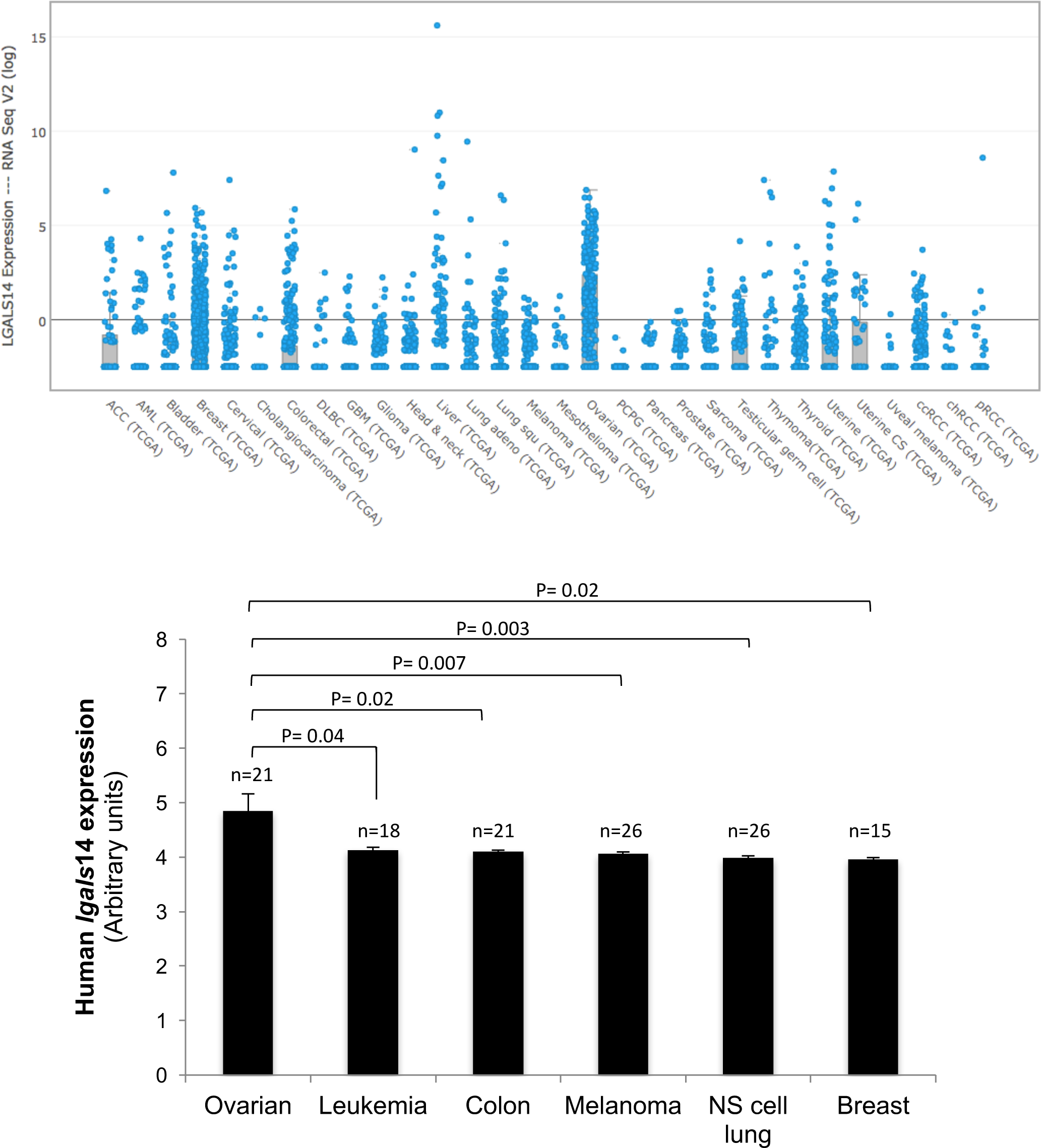
*lgals14* expression in cancer tissues and cell lines. (**A**) mRNA expression of *lgals14* from 53, 622 samples in 214 studies. (**B**) Human *lgals14* expression in the NCI-60 cancer cell lines.

### High expression of galectin-14 is associated with decreased survival of OC patients

Because epithelial ovarian cancer (EOC) is an aggressive type of cancer with limited treatment options, we focused our investigations on EOC, specially high grade serous ovarian adenocarcinoma (HGSA), the most aggressive subtype of EOC. Data mining using the Expression Atlas database, an open source that allows to study expression of specific genes across tissues or cell lines showed that *lgals14* was expressed in many EOC cells lines (Figure 3). Interestingly, EOC cell lines expressing the highest level of *lgals14* were derived from patients with HGSA. This finding included OVCAR-3, a common cell model to study EOC. An overall survival curve calculated by Kaplan-Meier method showed that high levels of *lgals14* were associated with a poorer survival (*p* = 0.016) (Figure 4). The expression of *lgals14* was also significantly higher in EOC tissues as compared to control (normal) tissues (*p* = 0.03) (Figure 5a). Furthermore, comparison between *lgals14* expression in EOC cases that are sensitive and resistant to treatment was very close to the threshold of significance (*p* = 0.1) (Figure 5b). Whether *lgals14* expression is driven by genetic alterations in EOC cell lines is a real possibility as *lgals14* had the highest percentage of genetic amplification in EOC when compared to other members of the galectin family, most notably *lgals1* and *lgals3* (Figure 6). Genetic alterations (mostly amplifications) of *lgals14* is indeed frequently observed in serous ovarian carcinoma and correlates with a poorer survival of patients with ovarian cancer (*Supplementary figure S1 and S2*).

**Figure 3:**
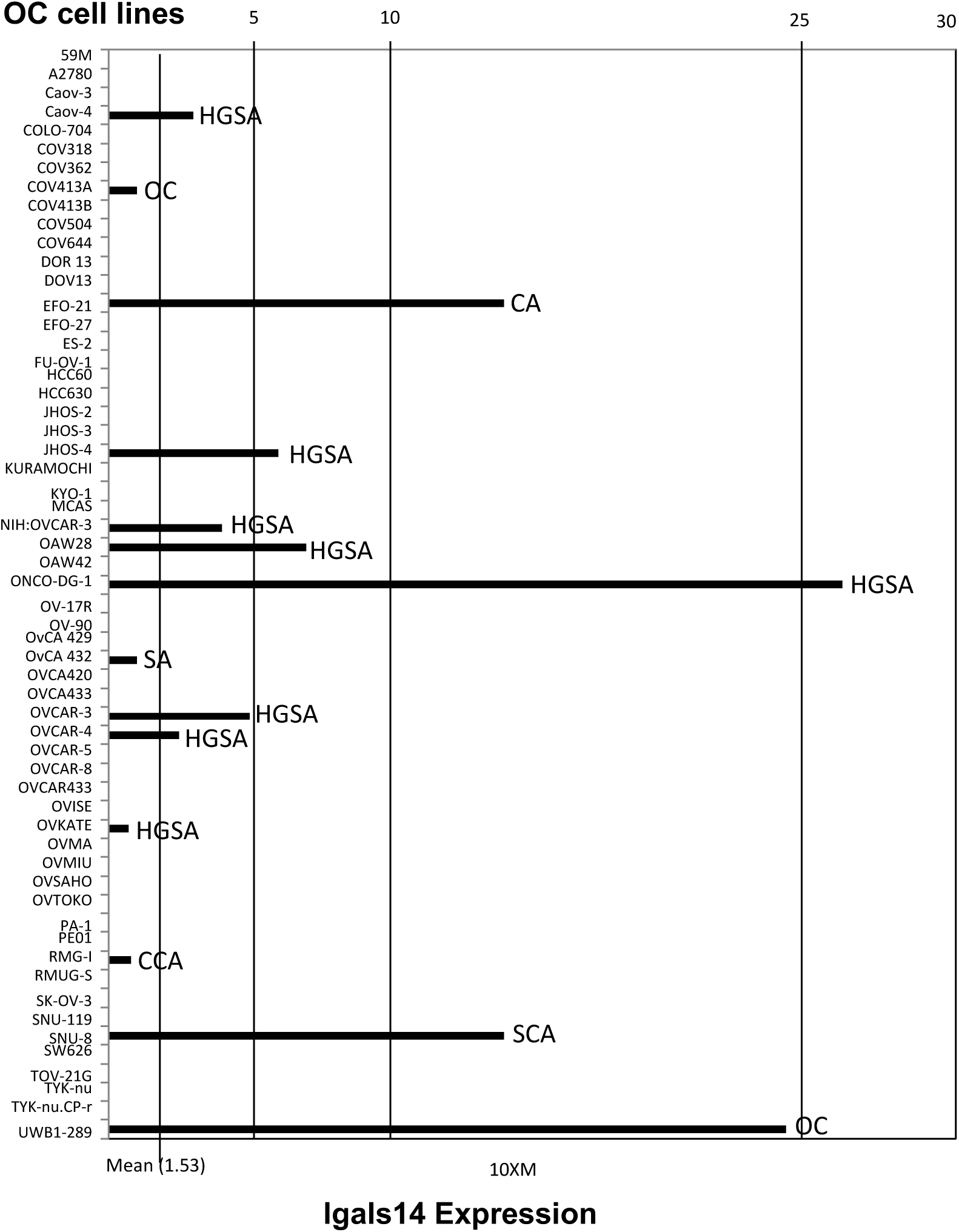
Expression of *lgals14* in human ovarian cancer cell lines. Data were obtained from the Expression Atlas database. HGSA; high-grade serous adenocarcinoma; OC; ovarian carcinoma; CA; ovarian cystadenocarcinoma; SA; ovarian serous adenocarcinoma; CCA; ovarian clear cell adenocarcinoma; SCA; ovarian serous cystadenocarcinoma.

**Figure 4:**
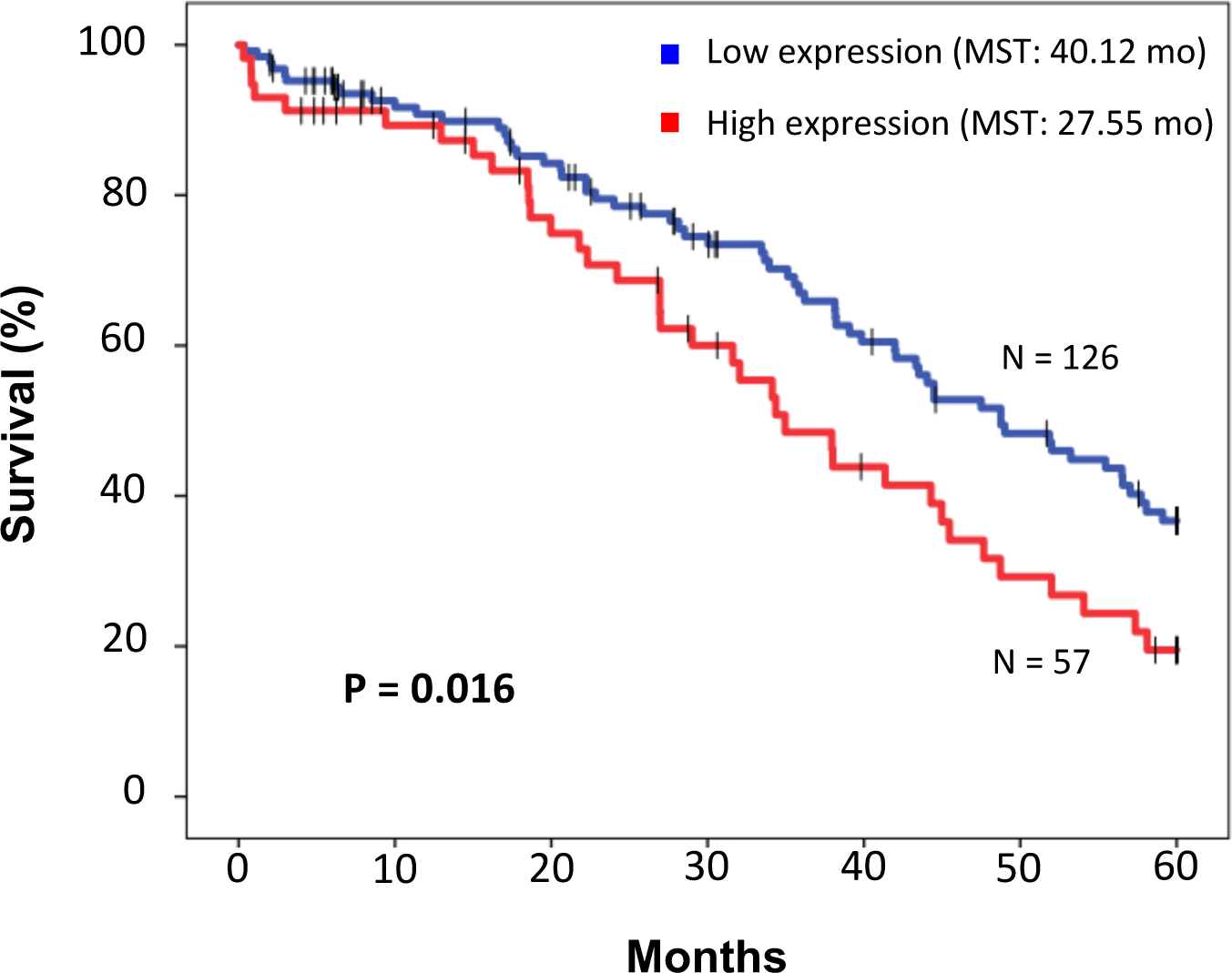
Kaplan-Meier survival analysis. Five-years OS of ovarian cancer patients with low or high expression of galectin-14 mRNA. Five-year overall survival (OS) of ovarian cancer patients according to *lgals14* expression levels. Data were obtained from the TCGA database.

**Figure 5:**
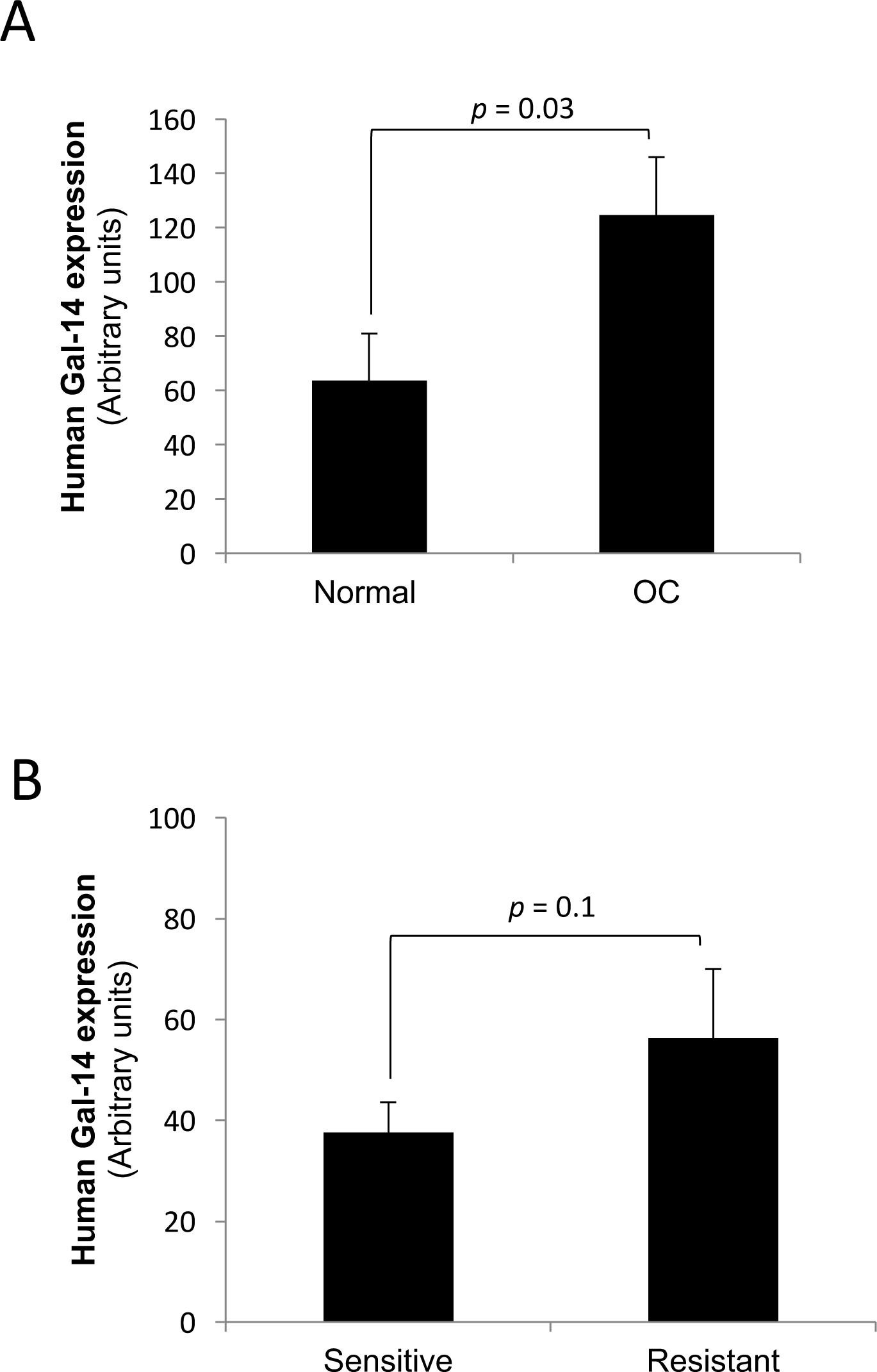
Expression of *lgals14* in ovarian cancer significantly higher than expression in normal ovarian epithelial cells. (**A**) Expression in normal and cancer ovarian epithelia. (**B**) Expression in high grade serous ovarian cancer according to resistance to platinum-based chemotherapy. Error bars represent SEM. All data were obtained from Geoprofile microarray datasets.

**Figure 6:**
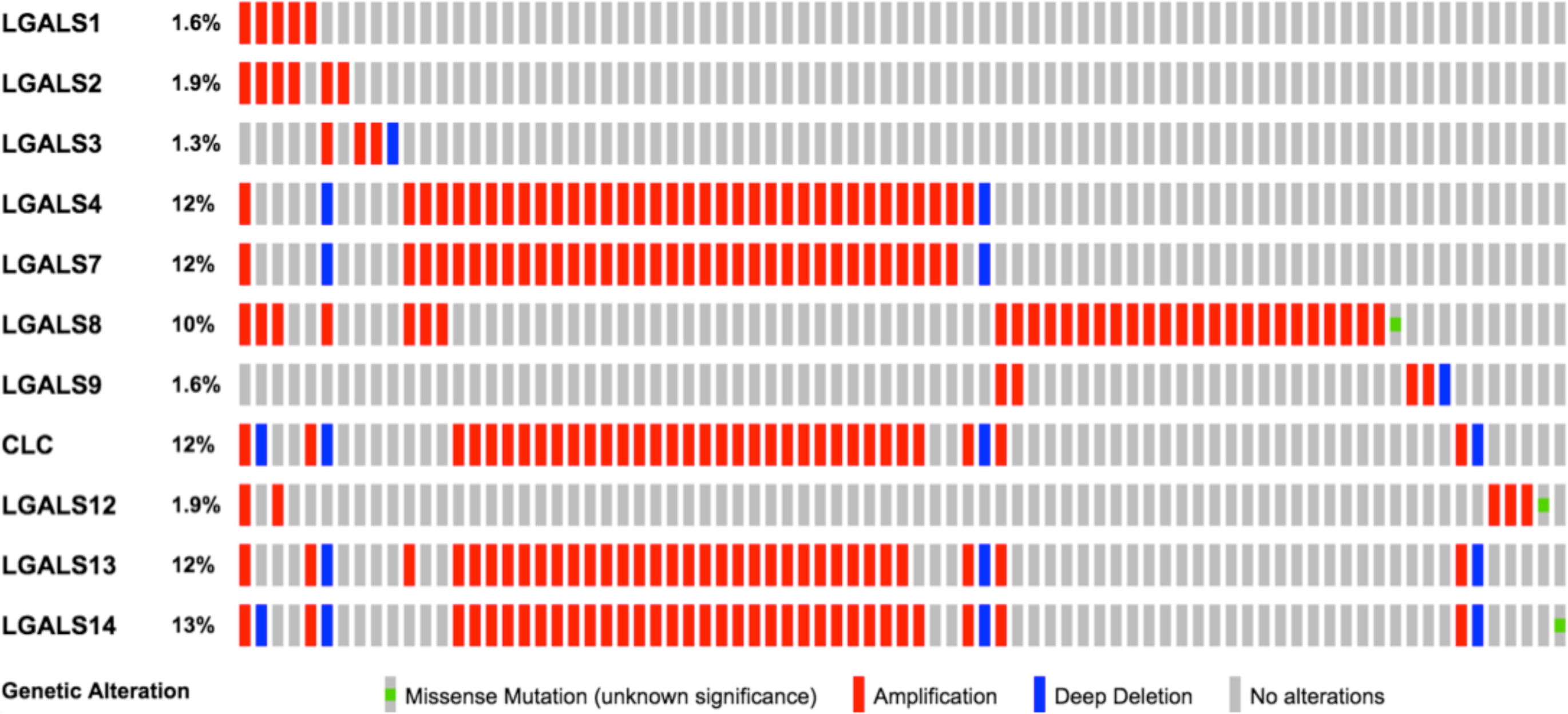
Genomic alterations in galectin genes in ovarian cancer. Galectins mutations across a set of ovarian serous cystadenocarcinoma cases.

### Validation of in silico data

We next carried out PCR and Western Blot analysis to validate the expression of gal-14 at the mRNA and protein levels. We also took the opportunity to look at other less well known galectin genes, including *lgals10, lgals12, lgals13*, and *lgals16*. We first confirmed that OVCAR-3 cells readily express high levels of *lgals14* (Figure 7a). No expression was found in HL-60 cell line, a common human leukemia cell model. *Lgals10* and *lgals16* were also expressed in OVCAR-3 cells, albeit at lower levels than *lgals14*. Expression of *lgals14* in EOC cell lines was also observed in TOV-1369 (TR), another HGSA cell line [21]. The TOV-1369(TR) cell line is a pre-chemotherapy cell line derived from a solid tumor (TOV) located in the fallopian tube (*Supplementary Table 1*). After ovarian cancer diagnosis, this patient received a treatment of paclitaxel and carboplatin for 5 months, two cycles of paclitaxel alone, 11 cycles of doxorubicin and a final treatment with Topotecan [21]. The OV-1369(R2) cell line is a metastatic cell line that was derived from the same patient at the end of the treatments [21]. Low but detectable expression of *lgals14* was also found in OV4453, another pre-chemotherapy HGSA cell line derived from ascites (OV) collected from a 70-year-old patient [22]. Both patients had a prior history of breast cancer. No detectable expression was observed in A2780, an ovarian endometroid adenocarcinoma cell line derived from an untreated patient, and SK-OV-3, a cell line derived from the ascites of a 64 year-old patient with an ovarian serous cystadenocarcinoma [23]. As expected, constitutive expression of *lgals14* was observed in placenta tissues. It was also found in the AML-14-3D10, an eosinophilic cell line derived from a patient with an acute myelogenous leukemia (AML) expressed *lgals14*, consistent with previous reports showing that *lgals14* is expressed in eosinophils, but not in other lymphoid or myeloid cells, including lymphocytes and macrophages [20, 24].

**Figure 7:**
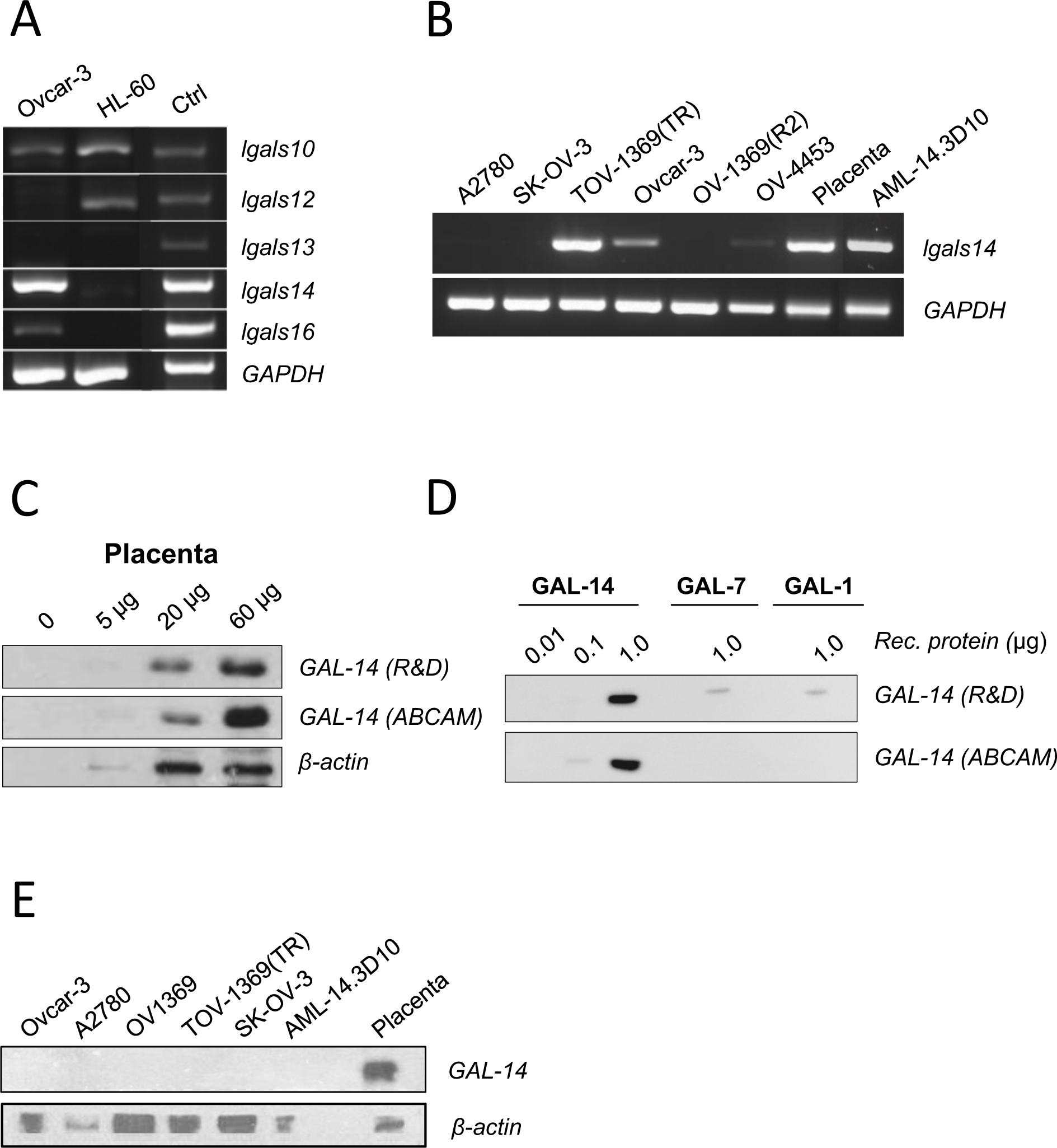
Galectin-14 expression in ovarian cancer cell lines. (**A**) mRNA expression of less well known galectins in OVCAR-3 and HL-60 cells. (**B**) The expression profile of *lgals14* in various EOC cell lines as measured by standard RT-PCR. DNA sequencing confirmed the identity of the amplicons (*Supplementary figure S3*) (**C**) Validation of gal-14-specific commercial antibodies by Western Blot. (**D**) Western Blot analysis of gal-14-specific antibodies against recombinant human gal-1, gal-7, and gal-14. (**E**) Galectin-14 protein expression profile in various EOC cell lines as measured by Western Blot.

### Galectin-14 protein levels

We next examined the expression of gal-14 at the protein level in EOC cell lines. First, we compared two commercially available anti-gal-14 antibodies using placenta and recombinant human gal-14. Our results showed that while both antibodies were capable of detecting gal-14 proteins, one of the antibodies showed low but detectable cross-reactivity with human gal-7 and gal-1 (Figure 7c, d). This antibody is a mouse monoclonal antibody (MAB5744, from R&D Systems) that was developed following immunization with human recombinant gal-14. The other antibody (ab150427, from Abcam) is a rabbit polyclonal antibody generated from an unknown (proprietary) peptide sequence. We thus used the Abcam antibody for the remaining of our experiments. We could not, however, detect gal-14 in protein extracts from ovarian cancer cell lines, nor with the AML-14.3D10 cells (Figure 7e). Our results obtained following transfection of a vector encoding human gal-14 in HEK-293 embryonic kidney cells, which do not normally express *lgals14* (Figure 8a), showed that we can detect gal-14 protein form in cellular extracts (Figure 8b). This suggests that while EOC and AML-14.3D10 cells express detectable levels of *lgas14*, they do not express detectable levels of gal-14 proteins.

**Figure 8:**
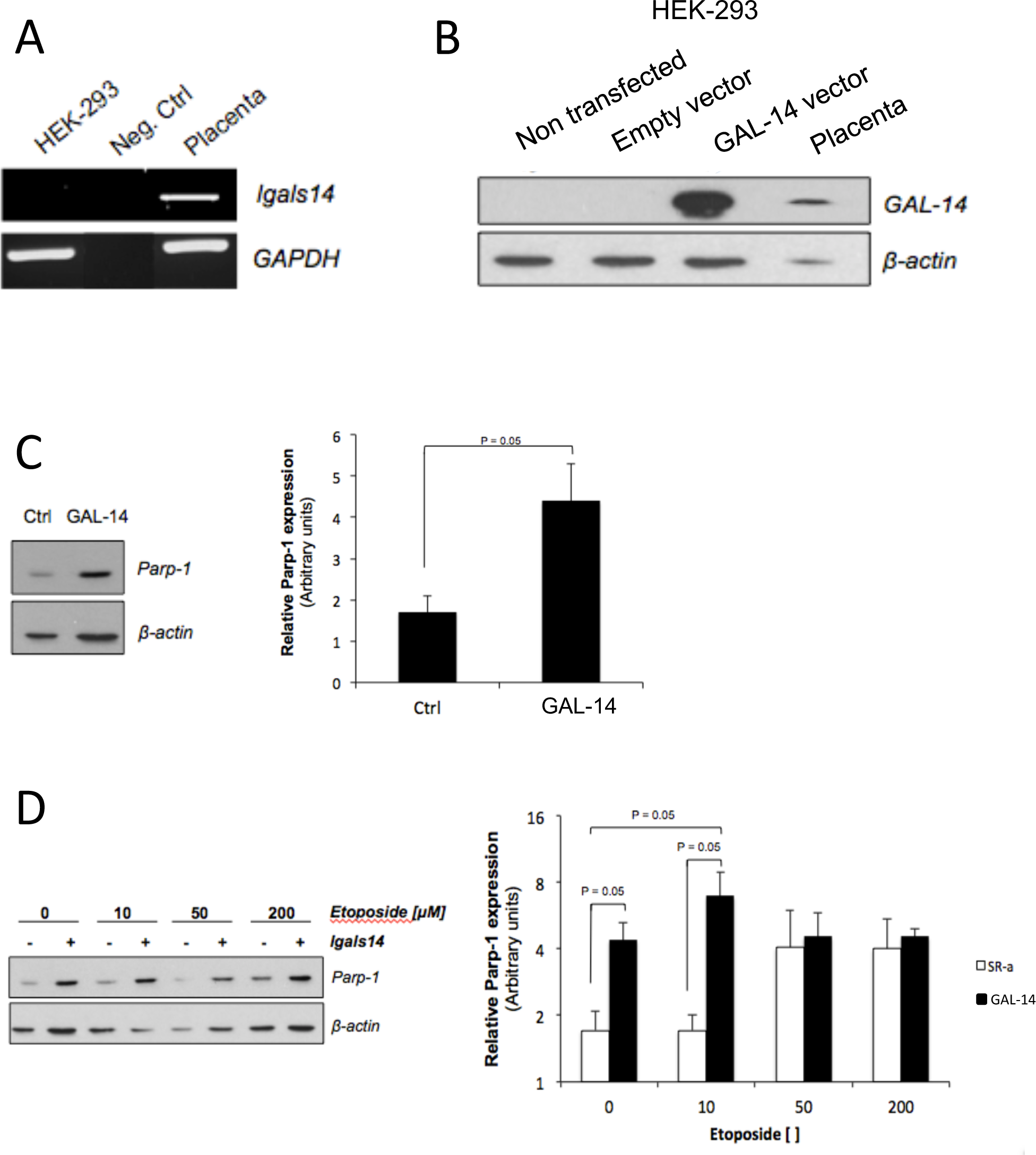
Parp-1 expression in HEK-293 transfectants. (**A**) RT-PCR analysis of *lgals14* in HEK-293 cells. (**B**) Validation by Western Blot of successful transfection of plasmid encoding *lgals14* in HEK-193 cells. Placenta was used as positive control. (**C**) Western Blot analysis of cleaved Parp-1 in transfected HEK-293 cells. (**D**) Parp-1 cleavage in transfected HEK-293 cells treated with increasing doses of Etoposide for 24h.

### Expression of gal-14 in HEK-293 cells

Our experiments with HEK-293 cells provided us the opportunity to explore, for the first time, the functional role of gal-14. Given the propensity of galectins to modulate apoptosis, we first compared PARP-1 cleavage in a (empty vector) control and cells transfected with a vector encoding human gal-14. Our results showed that *de novo* expression of gal-14 significantly (*p* < 0.05) increased the level of apoptosis in HEK-293 cells (Fig. 8c). It also increased apoptosis induced by low doses of etoposide, a common cancer chemotherapeutic agent (Fig. 8d). Not surprisingly, *de novo* expression of gal-14 in HEK-293 cells significantly reduced their *in vitro* growth (*Supplementary figure S4*).

## Discussion

In this study, using *in silico* and *in vitro* approaches, we have shed some light on the expression profile of gal-14 and started to explore its potential role in cancer. During our investigation, one type of cancer caught our attention: ovarian cancer. Our findings showed that *lgals14* expression is elevated in ovarian cancer cell lines derived from patients with HGSA, the most aggressive subtype of ovarian cancer. In addition, we observed that l*gals14* harbors more genomic alterations in patients with ovarian cancer than some of its famous family relatives, such as *lgals1* and *lgals3*. We also found that high expression of *lgals14* is associated with a shorter survival of patients with EOC, consistent with its preferential expression in HGSA. Additionally, we found that gal-14 expression in ovarian cancer cell lines is detectable at the mRNA level but not at the protein level. Our findings associated with HEK-293 transfectants suggest that this inability to detect gal-14 at the protein level is most likely due to the fact that these cells express very low levels of gal-14 proteins despite the fact that *lgals14* is readily detectable. This would be consistent with the our data showing that high expression of gal-14 increases basal levels of apoptosis. These results thus provide the first indications that gal-14 has a role of in cell death processes. Taken together, although the study of this protein is still in its infancy, we were able to provide novel insights into the expression patterns of this galectin and its involvement in cancer.

Our *in silico* study of *lgals14* expression in normal tissues agrees with previous studies showing that gal-14 is preferentially expressed in the placenta [16, 25]. To this day, the only data available about the expression of galectin-14 in other human normal tissues was published more than 15 years ago by researchers who studied gal-14 mRNA levels in multiple tissues by Northern Blot [17]. Yang’s team could not detect *lgals14* in other tissues. Our *in silico* study suggest, however, that gal-14 is possibly expressed at low levels in some tissues, most notably in adrenal and salivary glands, and possibly in the kidney and colon. Whether these low levels reflect expression in specific subpopulations within these tissues is a real possibility. Further studies using cell lines and tissue microarrays will be needed to address these issues.

Very few studies have examined the role of gal-14 in cancer. Using a gal-14-specific aptamer, Cho and colleagues have shown that gal-14 was expressed, albeit at low levels, in hepatocellular carcinoma cell lines, including SNU761, Huh-7, and SNU3058 [26]. Interestingly, the result of our *in silico* analysis of 53,622 DNA microarray samples from 214 cancer studies shows an elevated mRNA expression of *lgals14* in several cancers, including liver cancer. Yet, one interesting finding was that increased *lgals14* expression and mutation frequency of *lgals14* were clearly evident in tumors of the female reproductive tract, including breast, uterine and ovarian cancer, including the most malignant subtypes of ovarian cancer [27]. The involvement of galectins in female tract neoplasia has already been documented for some prototype galectins, such as gal-1 and −7. For example, increased expression of galectin-1 is found in uterine, breast, cervical, and ovarian cancers [28–32]. Gal-7 is also expressed at abnormally high levels in breast, ovarian and cervical cancer [12, 33–35]. When it comes the proteins homologous to gal-14, −10 and −13, the data is, again, very limited. The majority of studies have focused on the role of gal-10 in eosinophil-specific processes, and the functions of gal-13 in the maternal-fetal interface and in pregnancy-related events [36–39]. Therefore, our results are the first to give some insights into the expression of gal-14 in cancer, more specifically, in cancers of the female reproductive tract. In fact, our findings suggest that there might be an association between gal-14 and progression of EOC. The fact that *lgals14* is preferentially expressed in HGSA is particularly interesting given the limited treatment options for this subtype of cancer. Understanding the molecular mechanisms controling expression of gal-14 will also be needed to determine which pathways controls its expression in specific tumor subtypes. It is interesting to note that the promoter of *lgals14* contains consensus sites for both Nf-kB, Sp-1, and c/EBP, all of which have previoulsy been shown to control expression of gal-7 in cancer cells [33, 40].

Our results warrant further investigations on the role of gal-14 in EOC, most notably in its potential role in controlling cell death and the apparent discrepancy between the observations that higher *lgals14* expression is associated with lower survival of patients with EOC and the pro-apoptotic role of gal-14 observed in HEK-293 cells. One possibility is that gal-14’s pro-apoptotic function is cell type-specific. Another possibility is that the pro-apoptotic function of gal-14 in HEK-293 cells was due to the abnormally high levels of gal-14 following gene transfer, while EOC cells barely express the gal-14 protein, as we found in OEC cell lines, which keeps them protected from gal-14-induced apoptosis. This would explain why OV-1369(R2) cells, which were established following chemotherapy does not express gal-14 while TOV-1369(TR) (established prior to pre-chemotherapy) does. This would also explain why we were not able to detect gal-14 proteins in OVCAR-3 cells despite the presence of *lgals14*. This result was somewhat unexpected, given that transcription levels of a given gene often dictates the presence/absence of the corresponding protein. Yet, there are examples of galectins showing this type of molecular behavior. For instance, Satelli and colleagues showed that gal-1 and gal-3 proteins could not be detected in HEK-293 cells even though these cells expressed gal-1 and gal-3 mRNA [41]. Similar observations were also reported with gal-8 [41]. These findings support the existence of some type of translational repression mechanisms in ovarian cancer cells that keep gal-14 from being translated or specific proteolytic activity that effectively eliminates gal-14 as soon as it is synthesized.

It is becoming clear that members of the galectin family play an important role in cancer. Research over the past years has identified key roles for these lectins and their ligands in cancer immuno-editing and metastasis. We still need, however, to better understand the contribution of less well known galectins in cancer. This is particularly important for the development of anti-galectin drugs and for limiting their potential off-target effects. Overall, this preliminary study allowed us not only to provide novel insights into the expression patterns of this galectin in normal and cancer cells, but also to explore its potential role in cancer. It certainly sets the ground for future investigations of gal-14 in cancer, most notably in ovarian cancer. Although the study of gal-14 is still in its infancy, on a long-term basis, our results may contribute to the development of new and valuable anti-cancer strategies.

## Material and Methods

### In silico analysis

*In silico* analysis of galectin-14 expression was first conducted using datasets from the Gene Expression Omnibus (GEO) repository of the National Center for Biotechnology Information (NCBI) [42]. We examined microarray datasets obtained from profiling human normal tissues (GDS1096) [43], cancer cell lines (GDS4296) [44] and ovarian cancer cells (GDS3592 and GDS4950) [45, 46]. Additionally, other analysis were conducted using the public RNAseq datasets of large-scale cancer genomics projects obtained from the cBio Cancer Genomics Portal and the gene expression data across biological conditions from Expression Atlas database [47–49]. The putative transcription factor binding sites (TFBS) of galectin-14 gene sequence were identified using ALGGEN PROMO, a virtual laboratory for the study of TFBS in DNA sequences [50]. Analysis of public RNAseq datasets from the Cancer Genome Atlas (TCGA) were also conducted to correlate the expression of galectin-14 mRNA and the overall survival of patients with ovarian cancer.

### Cell lines, tissues, and reagents

The A2780 and OVCAR-3 ovarian cancer cell lines were kindly provided by Dr. Eric Asselin (Université du Québec à Trois-Rivières, QC, Canada), and the AML-14.3D10 eosinophil cell line was provided by Dr. Denis Girard (INRS-Institut Armand-Frappier, Laval, QC, Canada). These cells were maintained in RPMI 1640 medium supplemented with 10% [v/v] fetal bovine serum (FBS), 2 mM L-glutamine, 10 mmol/L HEPES buffer, and 1 mmol/L sodium pyruvate. The SK-OV-3 ovarian cancer cell line was grown in McCoy’s 5A medium supplemented with 10% [v/v] FBS, 2mM L-glutamine and 10 mM HEPES buffer at 37°C in a humidified atmosphere containing 5% CO_2_. The HEK-293 cell line was provided by Dr. Stéphane Lefrançois (INRS-Institut Armand-Frappier, Laval, QC, Canada) and it was maintained in Dulbecco’s modified Eagle complete medium (DMEM) supplemented with 10% [v/v] FBS. The high grade serous (HGS) ovarian carcinoma cell lines were a generous gift from Dr. Anne-Marie Mes-Masson (CRCHUM-Université de Montréal, Montreal, QC, Canada) and they were established from the ascites of a patient (OV-4453) and from a HGS ovarian tumor of a patient before (TOV-1369TR) and after (OV-1369 (R2)) chemotherapy treatment [21, 22]. These cell lines were maintained in ovarian surface epithelium (OSE) medium (Wisent, QC, Canada) supplemented with 10% [v/v] fetal bovine serum. All cell culture products were purchased from Life Technologies (Burlington, ON, Canada). The placenta tissue was provided by Dr. Cathy Vaillancourt (INRS-Institut Armand-Frappier, Laval, QC, Canada).

### RNA isolation andRT-PCR analysis

Total cellular and tissue RNA were isolated from cells using the TRIzol reagent (Life Technologies) according to the manufacturer’s instructions. First-strand cDNA was prepared from 2 µg of cellular RNA in a total reaction volume of 20 µL using the reverse transcriptase Omniscript (QIAGEN, Mississauga, ON, Canada). After reverse transcription, the cDNAs were amplified by PCR with genes specific primers (see Table 1) and using the following conditions: 94°C for 3 minutes, followed by 30 to 35 cycles of the following: 94°C for 60 sec, 62°C (*GAPDH*) and 65°C (l*gals14*) for 60 sec, and 72°C for 60 sec, followed by a final extension step at 72°C for 10 minutes. PCR was performed in a thermal cycler (MJ Research, Watertown, MA). The amplified products were analyzed by electrophoresis using 1.5% [w/v] agarose gels, SYBR Safe (Life Technologies) DNA gel staining and UV illumination.

### Western blot analysis

For whole cell extracts, cells were homogenized and resuspended in radioimmunoprecipitation assay (RIPA) lysis buffer (Thermo Fisher Scientific, Rockford, IL, USA) containing a cocktail of protease inhibitors (Roche, Laval, QC, Canada). For placenta extracts, the tissue was dissected, cut and homogenized in RIPA buffer with protease inhibitor, followed by centrifugation at 13,000 × *g* for 15 minutes at 4°C to pellet cell debris and collect lysate. Equal amounts of whole-cells and tissue extract (20-60 µg) were separated on 15% SDS-polyacrylamide gel and transferred onto nitrocellulose membranes (Bio-Rad Laboratories, Mississauga, ON, Canada). The membranes were first blocked with 5% milk [w/v] in TBS/0.05% Tween 20 [v/v] for 60 min at room temperature and subsequently blotted overnight at 4º C with primary antibodies (see Table 2). Secondary antibodies consisted of horseradish peroxidase-conjugated donkey anti-rabbit (GE Healthcare, Buckinghamshire, England), donkey anti-goat (R&D Systems) or sheep anti-mouse (GE Healthcare) IgG. Detection of immunoblots was performed by the enhanced chemiluminescence method using ECL detection reagent (GE Healthcare).

### Transfection and apoptosis assays

The vector encoding human Gal-14 gene (HG11371-CH) was obtained from Sino Biological (North Wales, PA, USA). The plasmid encoding human Gal-7 gene was obtained through the cloning of galectin-7 cDNA into sr⍺ eukaryotic expression vector using *Spe*I and *Bam*HI restriction enzymes as previously described (Campion et al. 2014). The controls were generated using the empty PCMV3 and Sr⍺ vectors. For transient transfection, the HEK-293 cells were plated at equal densities 24h prior to transfection and then transfected with the indicated vectors using the Lipofectamine 2000 reagent (Invitrogen) according to the manufacturer’s protocol. After 24h, transfected cells were treated with increasing concentrations of etoposide (0–200 µM) for 24h. Cells were lysed with 50 µL of RIPA buffer containing protease inhibitor at 4°C for 30 min and centrifuged at 13,000 × *g* for 15 min at 4°C. Equal amounts of protein were then separated and processed by Western Blotting following to the protocol previously described. Nitrocellulose blotting membranes were then probed with a rabbit anti-Poly-(ADP-ribose) polymerase (PARP)-1 polyclonal antibody (1:1000; Epitomics, Burlingame, CA, USA) and a mouse anti-β-actin antibody (1:10000; Sigma-Aldrich, St. Louis, MO, USA), followed by incubation with secondary antibodies and chemiluminescence detection.

### Statistics

Statistical significance of the experiments was evaluated using the unpaired Student’s t-test. Results were considered statistically significant at P≤ 0.05.

## Acknowledgments

The authors wish to thank Drs. E. Asselin and Denis Girard for providing cell lines. They also thank Ms. Marlène Fortier for her technical help.

## Supplementary data

**Figure S1:**
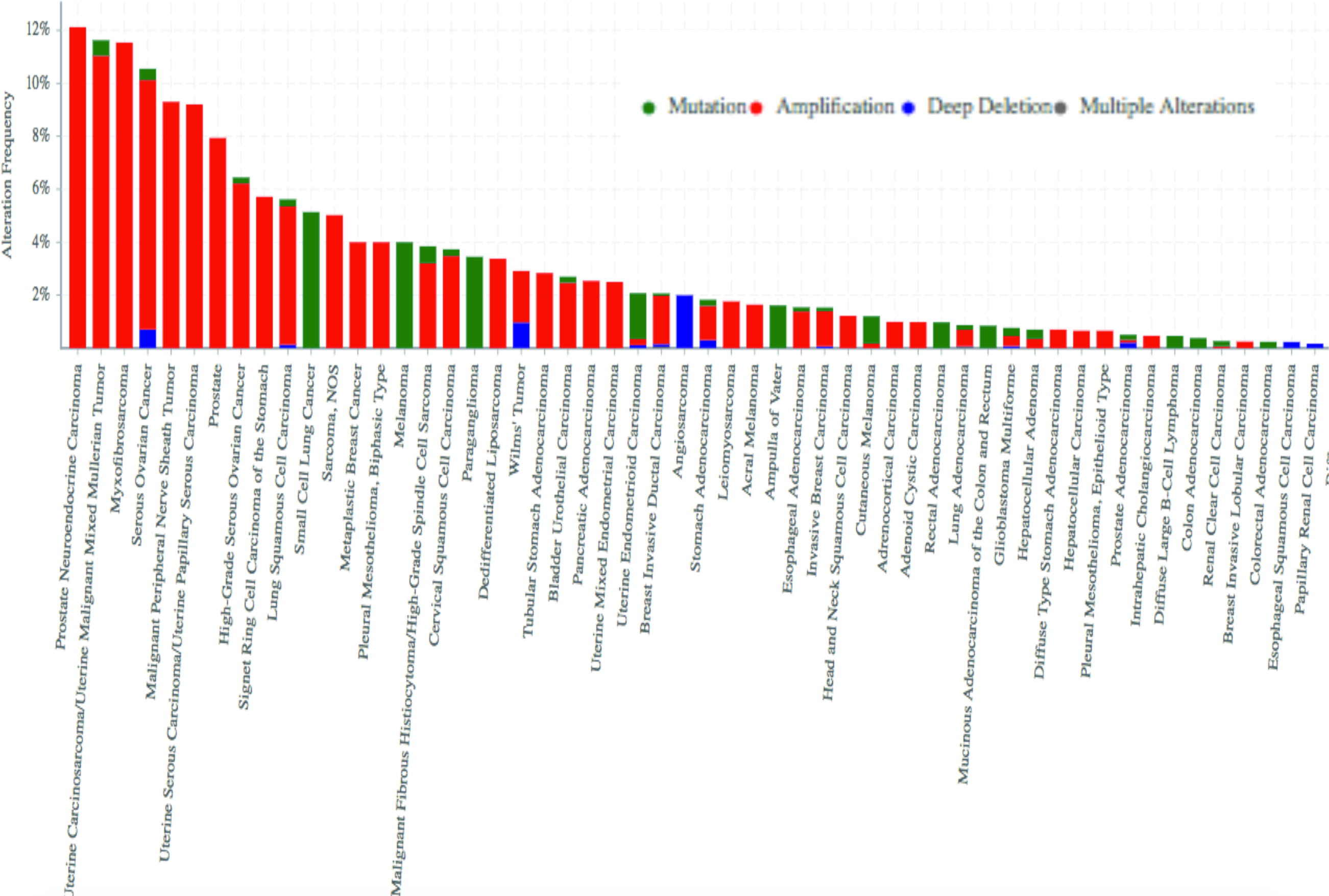
*lgals14* alterations in cancer. Data were obtained from the cBioportal database (cumulative data from 214 studies).

**Figure S2:**
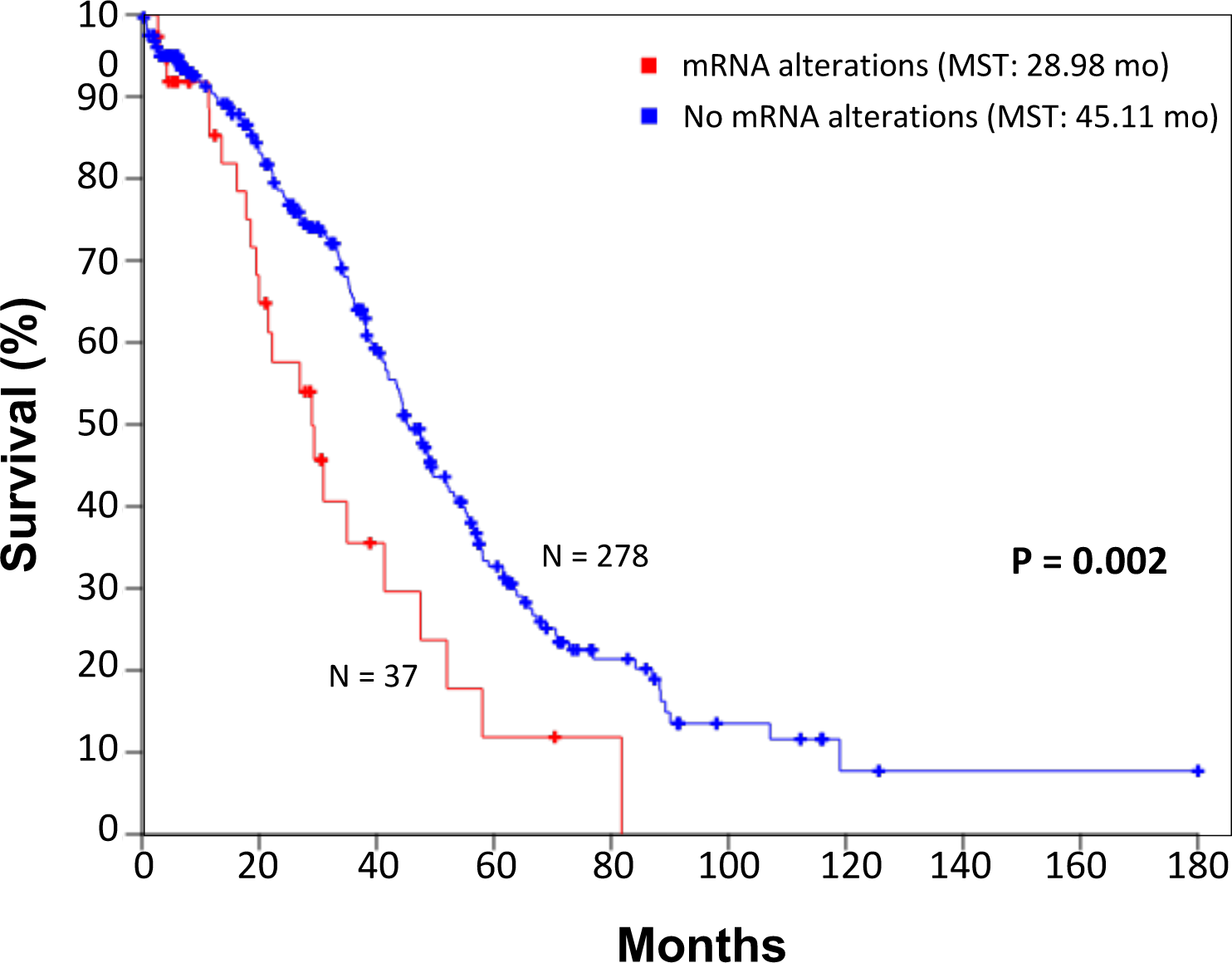
Overall survival of ovarian cancer patients according to the presence of genetic alterations in *lgals14*. Data were obtained through the TCGA database.

**Figure S3:**
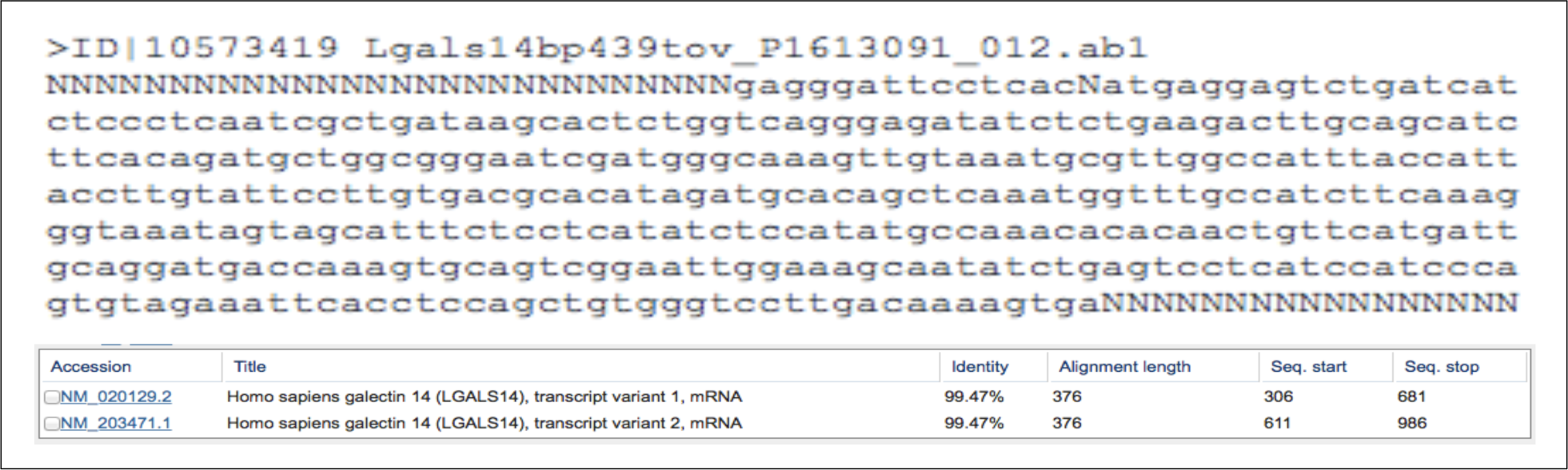
Validation of amplicons by sequencing using gal-14-specific primers.

**Figure S4:**
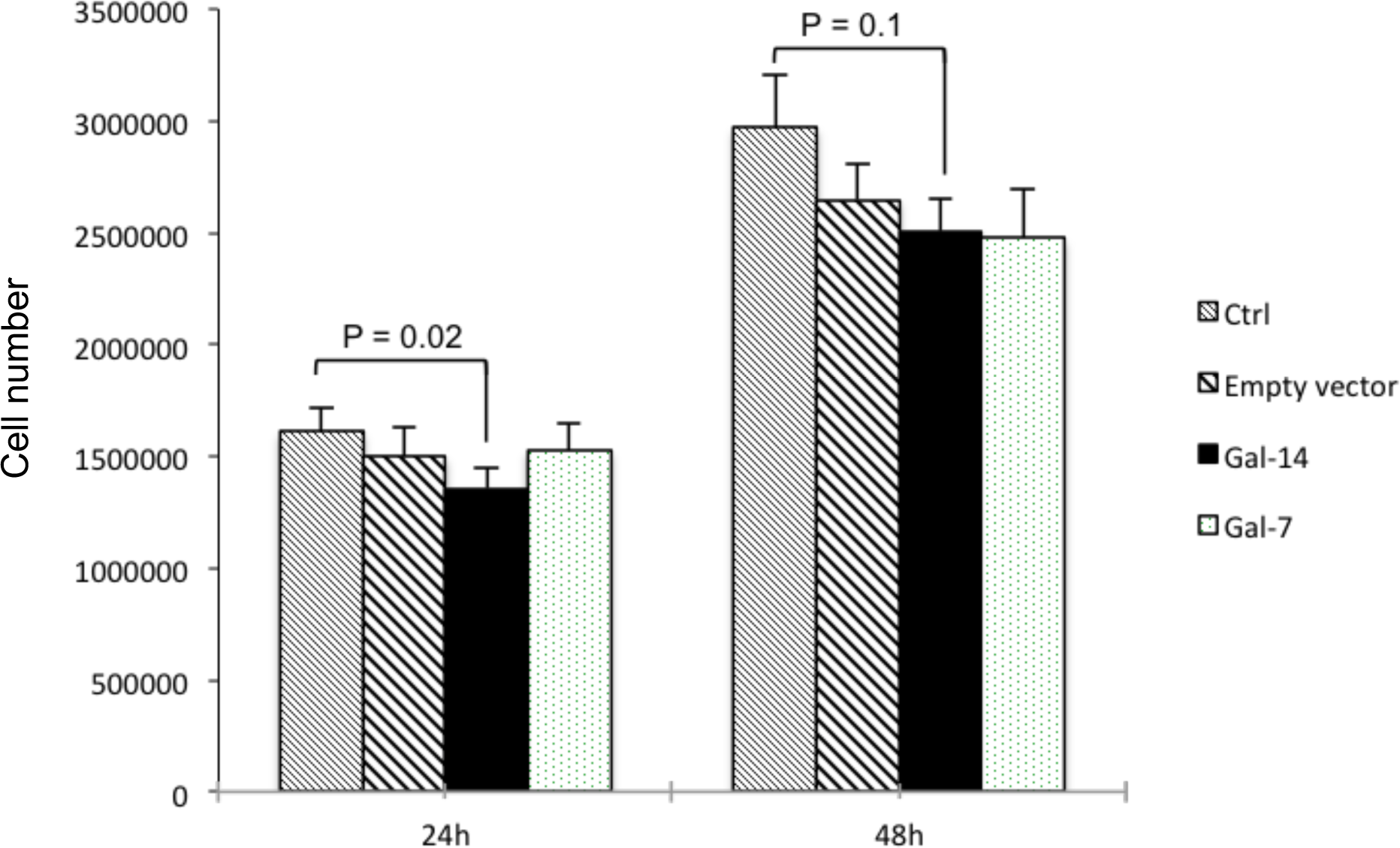
Cell proliferation following *de novo* expression of *lgals14*. HEK-293 cells were seeded into a 12-well-plate and counted at the indicated time using trypan-blue staining. The results are representative of four independent experiments.

**Supplementary Table 1.**

| Cell line | Origin | Subtype | Info |
| --- | --- | --- | --- |
| A2780 | Ovarian carcinoma | Serous | Established from tumour tissue from an untreated patient |
| Ovcar-3 | Adenocarcinoma | Serous | Established from the malignant ascites of a patient (Caucasian/ female/ 60 years) with progressive adenocarcinoma of the ovary. |
| SK-OV-3 | Adenocarcinoma | Serous | Derived from ascites from a 64-year-old patient. |
| TOV1369( TR) | Cystadenocarcinoma | High-grade serous papillary<br>Advanced stage (G3, IIIC) | Derived from fallopian tube from a 58-year-old patient. Pre-chemotherapy cell line. (TOV = solid/invasive tumor / TR = fallopian tubes). Patient with breast cancer + chemotherapy 14 months before |
| OV1369(2) | Cystadenocarcinoma | High-grade serous papillary<br>Advanced stage (G3, IIIC) | Post-chemotherapy cell line. OV = ascites<br>Patient had breast cancer + chemotherapy 14 months before |
| OV4453 | Adenocarcinoma | High grade serous | Derived from ascites from a 70-year-old patient. Previous history of breast cancer (5 years before). |

